# Transient Replication in Specialized Cells Favors Conjugative Transfer of a Selfish DNA Element

**DOI:** 10.1101/451864

**Authors:** François Delavat, Roxane Moritz, Jan Roelof van der Meer

## Abstract

Bacterial evolution is driven to a large extent by horizontal gene transfer (HGT) – the processes that distribute genetic material between species rather than by vertical descent. HGT is mostly mediated by an assortment of different selfish DNA elements, several of which have been characterized in great molecular detail. In contrast, very little is known on adaptive features optimizing horizontal fitness. By using single DNA molecule detection and time-lapse microscopy, we analyze here the fate of an integrative and conjugative element (ICE) in individual cells of the bacterium *Pseudomonas putida*. We uncover how the ICE excises and irregularly replicates, exclusively in a sub-set of specialized host cells. As postulated, ICE replication is dependent on its origin of transfer and its DNA relaxase. Rather than being required for ICE maintenance, however, we find that ICE replication serves more effective conjugation to recipient cells, providing selectable benefit to its horizontal transfer.

---

Integrative and Conjugative Elements (ICEs) are pervasive and permissive infestations of bacterial genomes^1–3^. ICEs display a dual lifestyle. Most cells in a population maintain the ICE chromosomally integrated, but under specific conditions a small proportion (estimated to between 1 in 10^2^–10^7^ cells depending on the ICE^3^) excises the ICE and produces an extrachromosomal ICE DNA-molecule^1–3^. The excised ICE-molecule can transfer into a recipient cell by conjugation, where it subsequently reintegrates. ICEs have attracted considerable interest because they frequently transfer and integrate into a wide taxonomic range of hosts, and carry gene functions of potential adaptive benefit to the host, such as genes coding for antibiotic or heavy metal resistance, plant symbiosis or xenobiotic compound metabolism^4,5^.

The model we use here is ICE*clc*, an element originally discovered in the soil bacterium *Pseudomonas knackmussii* B13, which bestows on its host a xenometabolic pathway to grow on the exotic compound 3-chlorobenzoate (3-CBA)^6–8^. Several characteristics of ICEs from the ICE*clc* family contribute to their remarkable ecological success in colonizing a large diversity of bacterial genomes^7^. In its integrated form, ICE*clc* is replicated with the host genome and remains largely without fitness cost on the host^9,10^. Although silent in exponentially growing cells, the ICE*clc* genes for horizontal transfer start to be expressed when all 3-CBA substrate in culture is depleted, turning some 3–5% of cells into a subset of specialized transfer competent (tc) cells (Fig. 1A)^11–13^. The ICE does not excise or transfer at this point, but only does so when tc cells are activated with fresh nutrients (Fig. 1A)^14^. Once turned into active donors, tc cells temporarily live before stalling cell division and lysing^12^. This fitness loss at population level is limited because of the small subpopulation size of tc cells, but these are highly effective in transferring ICE*clc*^14^. Beneficial for transfer is the induced formation by ICE*clc* of small tc cell groups that have an increased chance to contact recipients^15^.

**Fig. 1.**
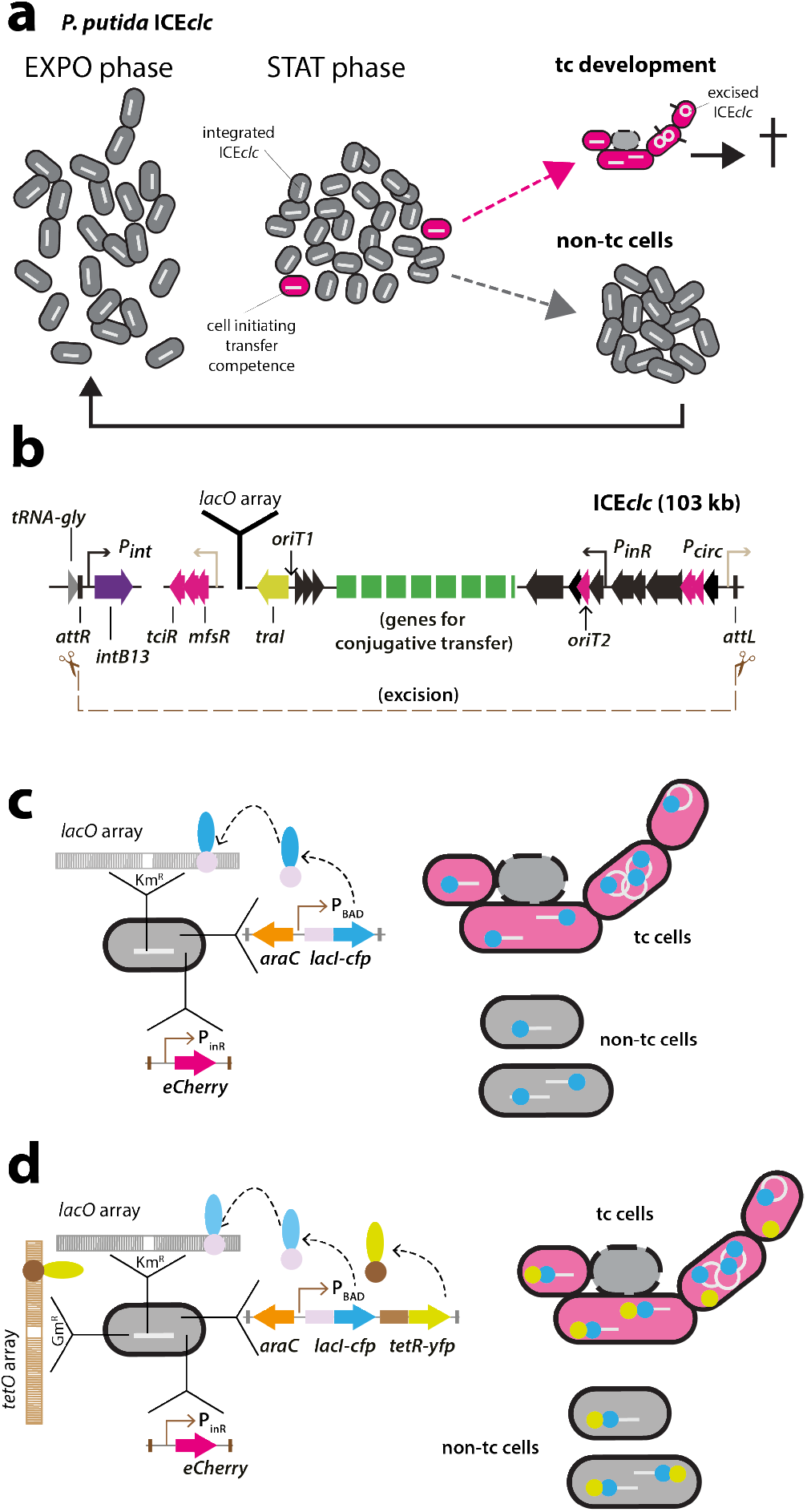
Principle of ICE*clc* detection in individual and transfer competent *Pseudomonas putida* cells. **a**, ICE*clc* remains integrated and silent in exponentially growing cells (EXPO, white bars in grey cells). 3–5% of cells in non-growing conditions (STAT) activate the core ICE promoters for its transfer competence program (magenta cells). Upon new nutrient addition, the transfer competent cells (tc) divide erratically, excising (white circle) and transferring the ICE (black protrusions from cells), and finally arresting cell growth and lysing (stippled cell outlines). Non-tc cells continue to divide normally in exponential phase. **b**, Schematic outline of ICE*clc*, its recombination boundaries, positions of genes mutated in this study and insertion position of the *lacO_ARRAY_*. **c, d**, LacI-CFP single or LacI-CFP/TetR-YFP double fluorescent foci formation by ectopic expression of a single copy chromosomally inserted arabinose-inducible *lacI-cfp*, or *lacI-cfp/tetR-yfp* gene construct. Fluorescent protein binding to the 240-fold copied cognate DNA binding site results in visible foci, as illustrated schematically. Cells are co-labeled with a single copy chromosomally inserted fusion of *echerry* to the P_inR_-promoter of ICE*clc*, which is active exclusively in tc cells.

Because of the limited tc-cell life span one would expect a number of specific adaptations favoring successful ICE transfer. A particularly critical moment for the ICE in the tc cell is when it is actually excising from the chromosome and engages in further conjugative steps. Indirect information from quantitative PCR in studies on other ICE models in *Bacillus subtilis* and *Vibrio cholerae* have suggested that ICEs can transiently replicate after excision, which has been regarded as a mechanism for its own maintenance and segregation among dividing daughter cells^16–18^. These studies, however, were population based and did not take individual cell fates into account. Since tc cells eventually disappear, producing multiple ICE copies may rather or also have been selected as a strategy to increase transfer chance. Previous evidence from single-cell studies have indeed suggested that individual donors can transfer to 2–3 surrounding recipient cells, the mechanism of which has remained elusive^14^. The main goal of the underlying study was thus to study replication of excised ICE*clc* in individual tc cells and to determine whether replication is advantageous for transfer fitness.

## Results

### ICE*clc*-DNA excision in individual tc cells

In order to differentiate and quantify single-copy integrated from excised ICE*clc*-DNA molecules in individual tc cells over time, we deployed the principle of fluorescent LacI-CFP fusion protein binding to a multi-copy integrated *lacO* array^19^. The *lacO_ARRAY_* was integrated on a single copy of the ICE*clc* in the genome of *Pseudomonas putida* (Fig. 1B, C). This strain was further tagged by a fluorescent reporter expressed uniquely in tc cells^20^. In addition, we constructed strains carrying an additional *tetO*-array nearby the ICE on the chromosome that can be bound by ectopically expressed fluorescent TetR-YFP (Fig. 1D). In normally replicating (non-tc) cells with integrated ICE*clc* we expected to observe 1–2 foci of LacI-CFP alone (when using the *lacO_ARRAY_* alone) or overlapping with TetR-YFP foci (when using cells with both integrated arrays). Upon ICE*clc* excision and consequent independent replication, we expected to see 3 or more fluorescent foci and potentially larger distances between TetR-YFP and LacI-CFP foci, exclusively in tc cells (Fig. 1C, D).

*P. putida* containing wild-type ICE*clc* tagged with the *lacO_ARRAY_*, ectopically expressing LacI-CFP showed a clear CFP focus in individual non-growing cells, but not when *lacI-cfp* was not induced (strain 5222, Fig. 2A). These cells are in non-dividing stage, and the single observed fluorescent focus is thus in agreement with a single chromosomal integrated ICE*clc* copy, formed by the attached LacI-CFP proteins to the *lacO_ARRAY_*. Foci were not visible in control *P. putida* strains with ICE*clc* but expressing only LacI-CFP, nor in *P. putida* with ICE*clc* and *lacO_ARRAY_* but without LacI-CFP (Fig. S1).

**Fig. 2.**
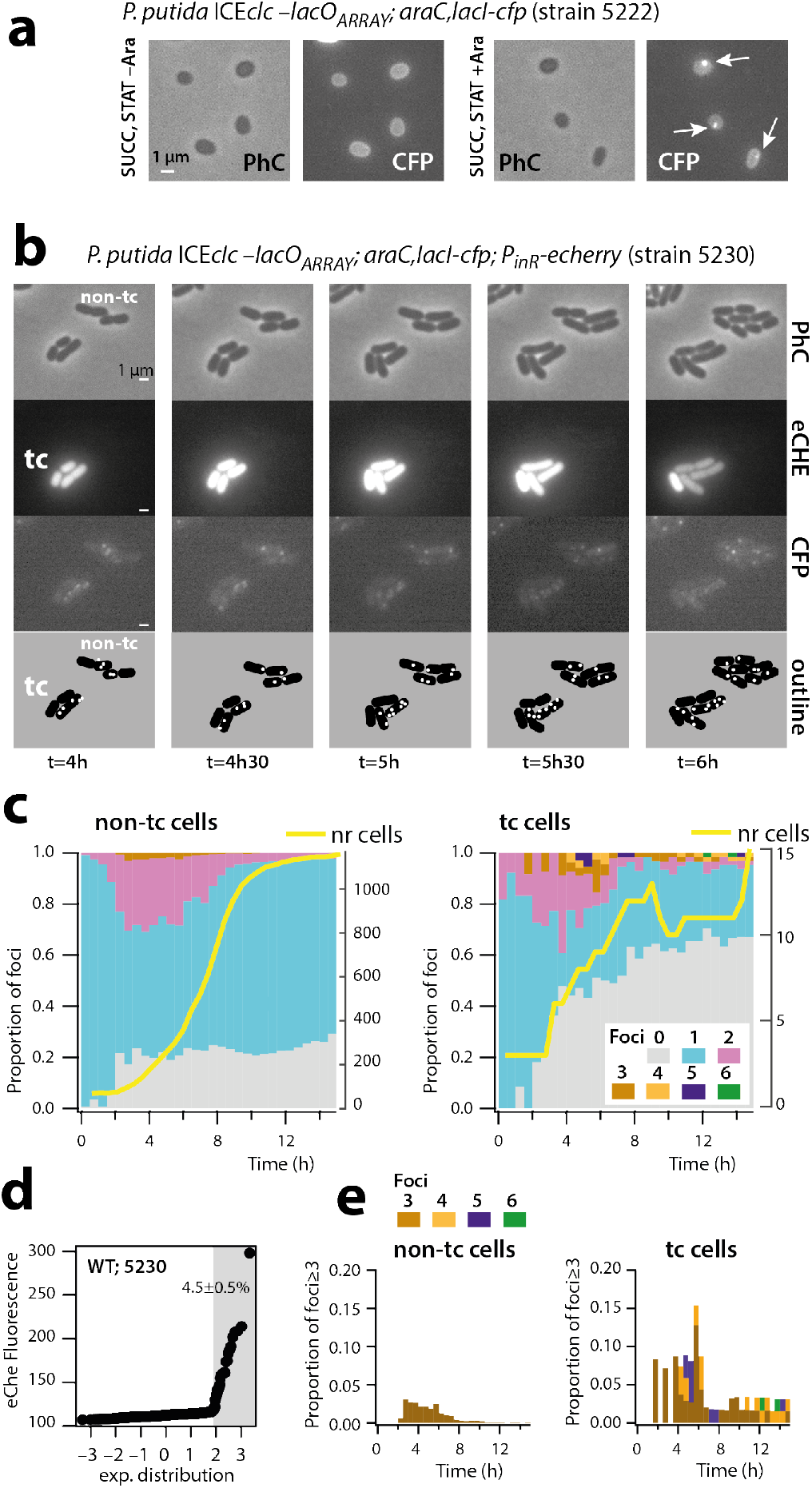
Increased LacI-CFP foci formation in transfer competent cells. **a-e**, CFP foci visible in *P. putida* stationary phase cells upon arabinose induction (**a**), and in tc and non-tc microcolonies growing on agarose mini disks (Note the stage of 6 foci in tc cell at t=5 h, **b**). Proportional foci distributions among growing non-tc and tc cells (nr cells, number of cells – right y-axis, **c**), with the initial discrimination of tc cells based on eCherry fluorescence (**d**), and foci distributions ≥3 among both cell types (**e**).

To measure ICE*clc* copy numbers and their temporal variation from fluorescent CFP foci in tc and non-tc cells, we deposited cells of a stationary phase 3-CBA–grown culture of *P. putida* ICE*clc–lacO_ARRAY_; araC,lacI-cfp;* P_inR_-*echerry* (strain 5230) on small agarose growth disks^14^. Cells grow exponentially to microcolonies as a result of included 3-CBA (Fig. 2B) and attain stationary phase after some 12 h (Fig. 2C). Importantly, because the seeding culture originates from stationary phase on 3-CBA, the population at the start of the experiment is composed of both tc and non-tc cells (Fig. 2B). These can be differentiated in the first time-lapse frame based on the eCherry fluorescence expressed from the chromosomally integrated P_inR_ reporter, which is active exclusively in tc cells^13,20^. On average, 4.5 ± 0.5% of individual cells in culture were representative for tc cells (Fig. 2D). It should be noted that the criterium of higher eCherry expression (as in Fig. 2D) is sufficient to classify cells into the tc cell category^21^, but not sufficiently exclusive to categorize cells as being non-tc, because some (true) tc cells may display low eCherry levels. For this reason, individual cells were further excluded from the non-tc class when their total number of offspring was less than eight. This criterium is based on the previously observed stalled cell division in tc cells^12^.

The majority of non-tc cells of *P. putida* strain 5230 displayed a single LacI-CFP focus (Fig. 2C, cyan-stacked bars). During population exponential growth (2-8 h after inoculation, yellow line in Fig. 2C, non-tc cells), some 20-30% of cells showed two foci (Fig. 2C, magenta-stacked bars). Two foci is in agreement with dividing cells replicating their chromosomal DNA, which, at some point, will thus have two ICE*clc* copies (producing two foci) before the chromosomes segregate among the daughter cells (see non-tc cell micrographs of Fig. 2B). A small proportion of cells (up to 4%) in the non-tc population displayed three CFP foci during exponential phase (Fig. 2E), which may be due to renewed replication forks, whereas up to 20% of cells had no detectable foci (Fig. 2C). Given that individual cells without detectable foci still divide on 3-CBA–containing media, we assume that such cells still contain ICE*clc* (note that the *clc* genes on the ICE are necessary for growth on 3-CBA^8^). Absence of visible foci in such cells is therefore more likely either an imaging artefact (e.g., the CFP-spot being out of image focus in the cell), or caused by a variability in induction by externally added arabinose.

In contrast, and although their overall number was much lower, tc cells showed a very distinct foci pattern from non-tc cells (Fig. 2C, E; tc cells). During their division phase, some 20% of tc cells displayed two CFP foci, but two foci were detected in cells even before the onset and after the end of population growth (Fig. 2C). A much larger proportion of tc cells showed no CFP foci at all, whereas in strong contrast to non-tc cells, up to 15% of tc cells displayed three and up to six CFP foci (Fig. 2B, 2E). The microcolony shown in Fig. 2B further illustrates the dynamic appearance of foci in tc cells. The foci distributions between tc and non-tc cells were highly significantly different in a Fisher’s exact test (p=0.0005). The consistent higher number of LacI-CFP foci in tc than in non-tc cells and the fact of having three or more foci in individual cells suggested the ICE to have excised and undergoing replication of the excised form. The much larger proportion of tc cells without any visible fluorescent focus compared to non-tc cells (60 vs. 25%, Fig. 2C) appearing over time, might be due to cells losing the ICE (postulated in Ref^14^, although the tc cell example of Fig. 2B shows a dividing tc cell without CFP foci). In addition, as for non-tc cells, imaging artefacts or variability of arabinose uptake, which is needed for expression of LacI-CFP, may have led to cells without visible CFP foci.

To further confirm ICE*clc* excision we used a *P. putida* derivative (strain 5601) containing, in addition to the *lacO_ARRAY_* on the ICE itself, a *tetO_ARRAY_* integrated in the Pp_1867 locus 12 kb upstream of the position of ICE*clc attR* on the *P. putida* chromosome (Fig. 1D). A diagram of TetR-YFP foci positions plotted as function of the longitudinal cell axis size illustrates the ongoing chromosome replication in non-tc and tc cells (Fig. 3). Cells with two observable YFP foci tend to have lengths of 1.8 µm upwards up to a length of 2.5–3.0 µm, after which the two daughter cells separate (Fig. 3A). Longer non-tc cells tend to display proportionally larger TetR-YFP interfocal distances, with positions symmetrical to the cell middle, indicative for segregating replicating chromosomes (Fig. 3C, compare red and blue dots). The majority of non-tc cells (90.4%) with a single YFP focus are smaller (0.8–1.8 µm), and the relative distance of that focus to the cell middle is less than 0.5 µm (Fig. 3A, light brown dots). An estimated 89.1% of non-tc cells with a single CFP focus were sized in between 0.8–1.8 µm, whereas 10.9% were larger than 1.8 µm and thus may have carried two CFP foci, although only one was visible. Replicating chromosomes were less clearly visible for CFP than for YFP foci in cells larger than 1.8 µm (Fig. 3A). The average distance between CFP and YFP foci in non-tc cells with one visible focus of each (i.e., cell size range 0.8–1.8 µm) or in cells with sizes in between 1.8–2.5 µm with two visible foci of each (in the same replichore) was close and not significantly different (199 ± 126 nm vs 172 ± 111 nm, p=0.3032 in ANOVA, Fig. 3B). This is indicative for integrated ICE*clc*, with closely juxtapositioned LacI-CFP and TetR-YFP binding sites (Fig. 3C).

**Fig. 3.**
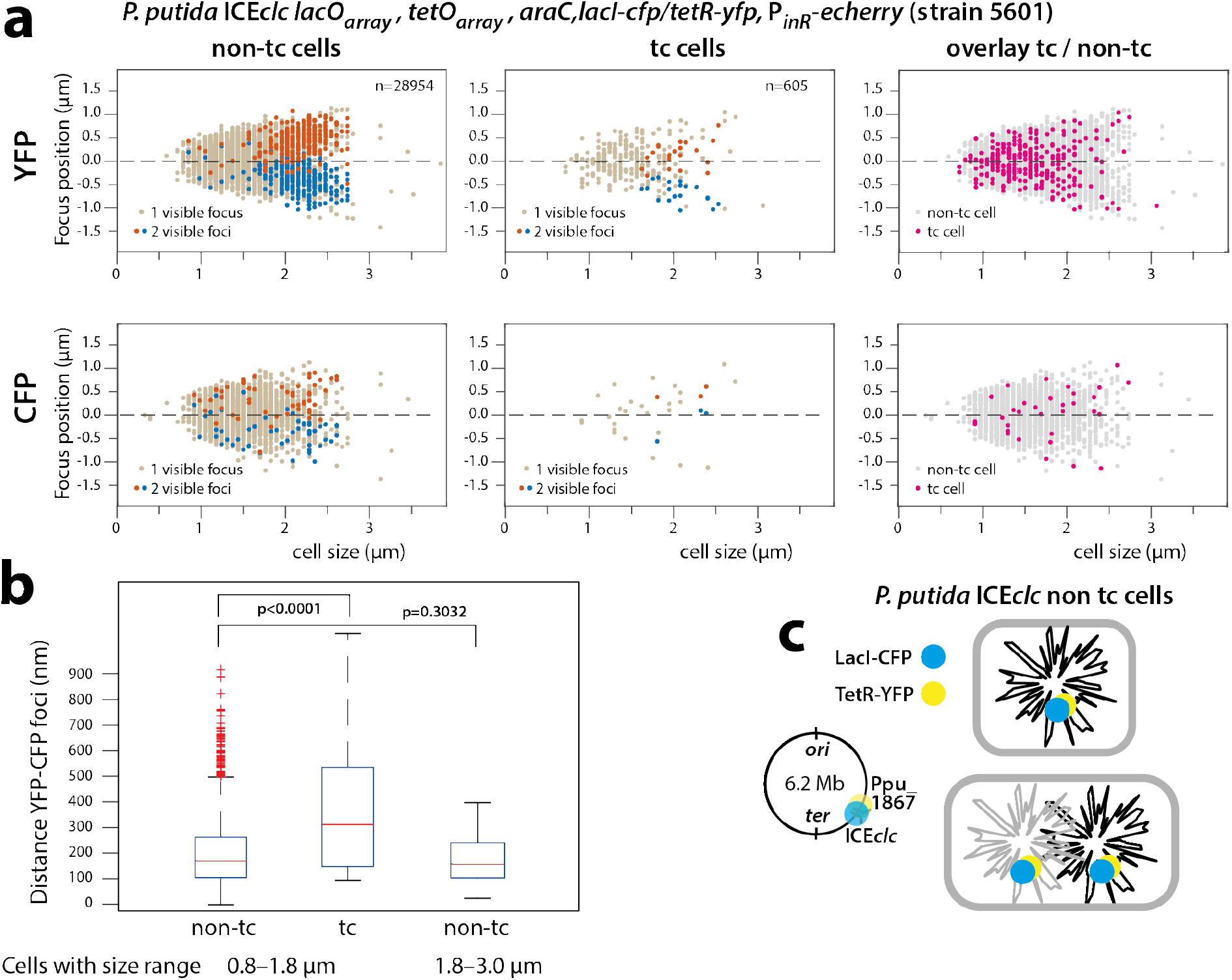
Excision of the ICE in *P. putida* tc cells co-labeled with LacI-CFP and TetR-YFP. **a**, Foci positions in individual non-tc or tc cells plotted as a function of cell size. Note visible chromosome replication (double TetR-YFP foci) in cells > 1.8 µm and note how TetR-YFP foci in tc cells follow the general positional pattern of non-tc cells (overlay). CFP foci in general less abundant and visible in co-labeled strains. n, number of analyzed cells in each category. **b**, Increased interfocal (TetR-YFP/LacI-CFP) distances in small tc compared to non-tc cells (0.8–1.8 µm) with on average a single chromosome copy (p<0.0001 in ANOVA), but not between smaller and larger non-tc cells with two foci pairs (partial chromosome replication, p=0.3032). **c**, Because of the position of the integrated *IacO*- and *tetO_ARRAY_* on the *P. putida* chromosome, it is unlikely that four pairs of foci would be visible as a result of renewed chromosome replication before daughter cell separation.

tc cells ranged similar in size as non-tc cells (Fig. 3A) and YFP foci positioned in broadly the same trends (Fig. 3A, YFP overlay). The distance between CFP and YFP foci positions in tc cells of 0.8–1.8 µm (which on average have a single chromosome), however, was on average twice as large as in non-tc cells (385 ± 287 nm, p<0.0001 in ANOVA). This indicates that ICE*clc* and the nearby chromosome locus are physically separated, which is in agreement with the hypothesis of ICE*clc* being excised in tc cells. The number of tc cells in the 1.8–2.5 µm category with two YFP–CFP foci couples was insufficient for statistical comparison.

### ICE-factor dependent replication of excised ICE

In order to determine whether the observed multiple ICE copies in tc cells (3–6) were the result of ICE replication, we quantified temporal variations in CFP foci in a variety of ICE-mutant strains of *P. putida*. In *P. putida* with an ICE*clc* deleted for the regulatory gene *mfsR* (Fig. 1B), equipped with the *lacO_ARRAY_* and the inducible *lacI-cfp* system (strain 5233), the proportion of tc cells in stationary phase increased to 45.4 ± 6.3% (Fig. 4A)^22^. *P. putida* ICE*clc-∆mfsR* (strain 5233) cells showed some overt displays of multi LacI-CFP foci in individual tc cells (Fig. 4B). This example is also illustrative for the dynamic movement of the various LacI-CFP foci over time in individual non-dividing tc cells, suggesting some active mechanism for their redistribution (see Movie S1). Dividing non-tc cells of this ICE-hyperactive strain 5233 with deleted *mfsR* still mostly displayed one or two LacI-CFP foci, with small proportions of cells showing 3 foci (Fig. 4C, left). In contrast, tc cells carried significantly higher proportions of 3, 4, and up to 6 foci than non-tc cells (Fig. 4C, right, p-value in Fisher’s exact test: 0.0167). It should be noted that, given the large number of tc and non-tc cells, we did not manually inspect the correctness of automated segregation (in contrast to data shown in Fig. 2 when using strain 5230 with wild-type ICE). This may have resulted in some faulty segregated non-tc “cells” carrying more than 3 foci, which actually were double cells. Although the *∆mfsR* deletion yields a much larger proportion of tc cells, this mutation does not impair the assumed replication of ICE*clc*.

**Fig. 4.**
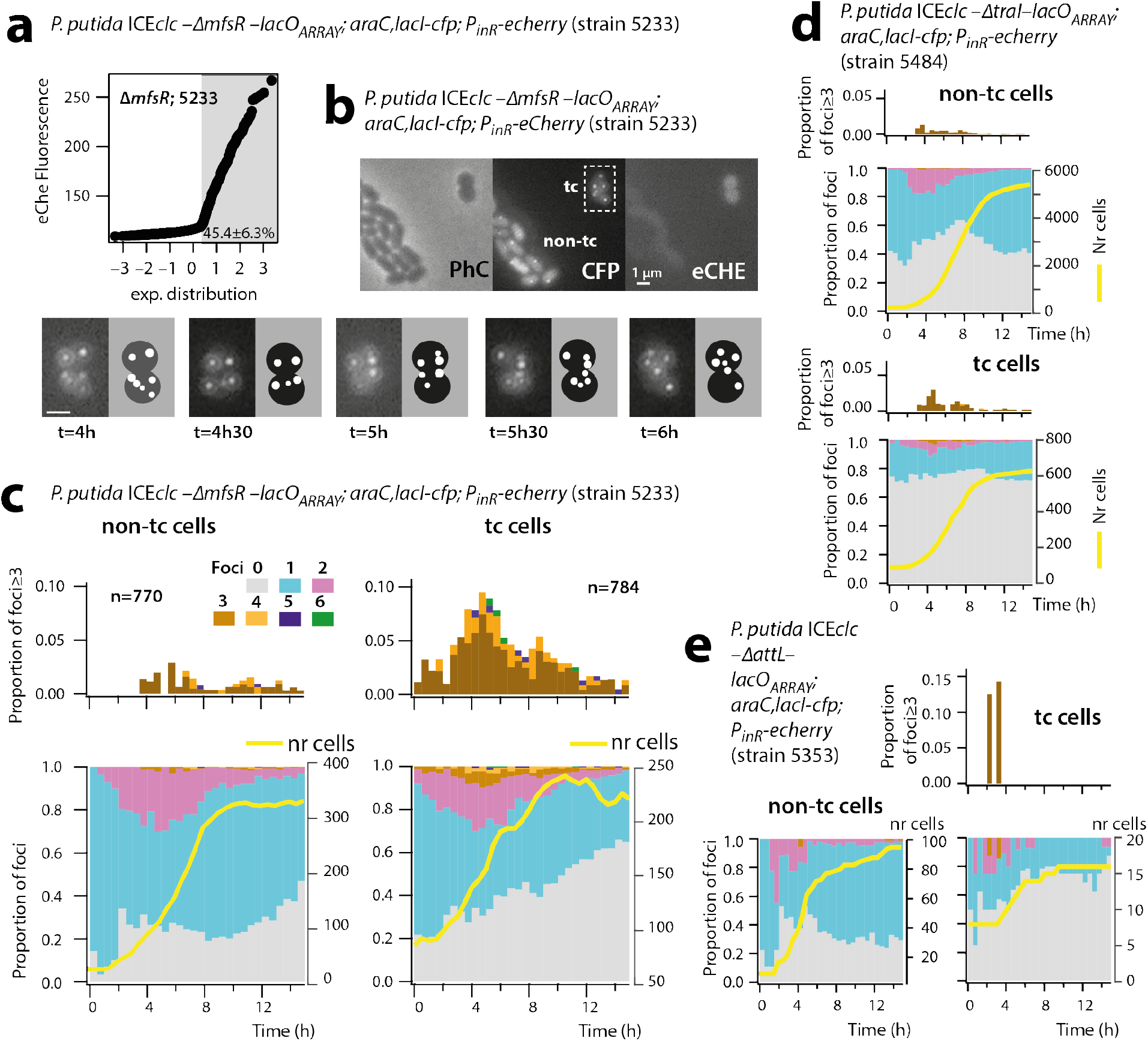
Effect of ICE mutations on tc cell foci distributions. **a-c**, Deletion of the *mfsR* global regulator increases the proportion of tc cells to 50% in stationary phase (**a**), with tc cells displaying multiple LacI-CFP foci (**b**). Note the slightly more roundish shape of the tc cells. **c**, Proportions of cells with ≥3 LacI-CFP foci in tc versus non-tc cells. **d, e**, Mutations inactivating the *traI* relaxase gene (**d**) or deleting the *attL* recombination region, disabling excision (**e**), result in fewer foci in tc cells.

The proportion of tc cells of a *P. putida* strain carrying an ICE in which the attL excision-recombination region was deleted, but otherwise with similar *lacO_ARRAY_*, *lacI-cfp* and *echerry* labels (strain 5353) was similar as for ICE wild-type (4.4±1%). However, except for a few sporadic time points, both non-tc and tc cells of strain 5353 did not display more than two foci (Fig. 4D). This is in agreement with our hypothesis that the ICE cannot excise in this mutant and therefore, that LacI-CFP foci solely indicate chromosomally integrated ICE copies. The tc cell proportion was higher in a *P. putida* strain carrying ICE*clc* with a deletion in the traI gene (Fig. 1B, 13.1–15.7%), which in other ICE-systems has been implicated in replication of the excised ICE^17,23^. In this strain (*P. putida* 5484) the proportion of tc cells displaying three and a few instances of four foci was clearly lower than in strain 5230 with wild-type ICE, (compare Fig. 4D and 2E), whereas those of non-tc cells were similar. This indicated that tc-cell LacI-CFP foci numbers higher than 3 are indeed the result of a replicative process that involves the TraI relaxase. Finally, foci numbers in tc cells of *P. putida* containing an ICE*clc* with a deletion in *oriT1*, one of the origins of transfer on which the TraI relaxase is acting^24^, never surpassed a maximum of three (Table 1). In contrast, *P. putida* with a deletion in the alternative origin of transfer oriT2 showed three percent of tc cells with 4 foci, which was more similar to wild-type (Table 1). This indicates that the *oriT1* region is important for the temporary replication of ICE*clc* upon excision.

**Table 1.** Effect of *oriT* mutations on the normalized LacI-CFP foci distributions in differentiated transfer competent cells

| *P. putida* strain | Cell type | Total N° cells | Percent of cells with n° foci |  |  |  |  |  |
| --- | --- | --- | --- | --- | --- | --- | --- | --- |
|  |  |  | 1 | 2 | 3 | 4 | 5 | 6 |
| 5230 (ICE*clc* wt) | tc | 77 | 49 | 38 | 6 | 4 | 1 | 1 |
|  | non-tc | 1904 | 59 | 38 | 3 | 0 | 0 | 0 |
| 5712 Δ*oriT1* | tc | 94 | 86 | 12 | 2 | 0 | 0 | 0 |
|  | non-tc | 4206 | 85 | 15 | 0.07 | 0 | 0 | 0 |
| 5713 Δ*oriT2* | tc | 149 | 73 | 20 | 6 | 3 | 0 | 0 |
|  | non-tc | 3581 | 76 | 23 | 2 | 0 | 0 | 0 |

Calculated p-values in Fisher's exact test of tc foci distribution comparisons, alternative hypothesis "two-sided" : 5230 vs 5712 p=0.0005; 5230 vs 5713 p= 0.0125; 5712 vs 5713 p=0.0490.

### Cells with more excised ICE*clc* copies transfer more frequently

To test the potential relation between ICE copy numbers in tc cells and success of ICE horizontal transfer, we mixed donors of *P. putida* ICE*clc-∆mfsR* tagged with *lacO_ARRAY_* and LacI-CFP with a conditional fluorescent *P. putida* recipient strain as bait. The recipient strain (strain 5248) fuses a promoterless *echerry* gene downstream of the *attB* recombination site^14^ and additionally expresses LacI-CFP. Integration of ICE*clc* into the conditional trap results in placement of the constitutive outward-facing P_circ_ promoter (Fig. 1B) directly upstream of *echerry*. Even though this strain captures only ~20% of all integration events (the others going into alternative *attB* sites on the chromosome formed by genes for tRNA-glycine^25^), one can quantify the numbers of foci in donor cells appearing in contact with eCherry-forming recipients, and compare their LacI-CFP foci distribution to that observed in all tc donor cells without recipient. Figure 5 shows two distinct examples of such transfer events. In figure 5A, the tc donor cell displays 4-5 LacI-CFP foci (visible after 1.5–3 h), leading to an mCherry-producing transconjugant visible at t=21 h. In figure 5B one can see how the incoming ICE*clc* is bound by the recipient‘s LacI-CFP (shown at t=14 h in two cells neighbouring the tc donor cell). One of those disappears over time, possibly as a result of aborted replication and lack of integration (cell labeled with ‘a’ in Fig. 5B, t=14 h). The other LacI-CFP remains and eventually leads to a recipient producing eCherry (at t=21.5 h), indicative for its proper integration into the conditional trap (Fig. 5B, the full time series of both events is shown in Figure S2). On average, we found a time span of 3.5–10 h in between the appearance of a LacI-CFP recipient focus (indicative of ICE transfer to the recipient) and detectable mCherry expression (indicative of ICE integration in the recipient‘s genome). Across 20 detected transfer events with ICE*clc* integrated in the recipient’s conditional trap, the identified tc cell donors displayed more LacI-CFP foci than expected from the foci distribution seen for tc cells in general (Fig. 5C, p-value=0.0004998 in Fisher’s exact test comparing foci distributions). This is thus a strong indication that donors with multiple ICE copies preferentially contribute to ICE transfer.

**Fig. 5.**
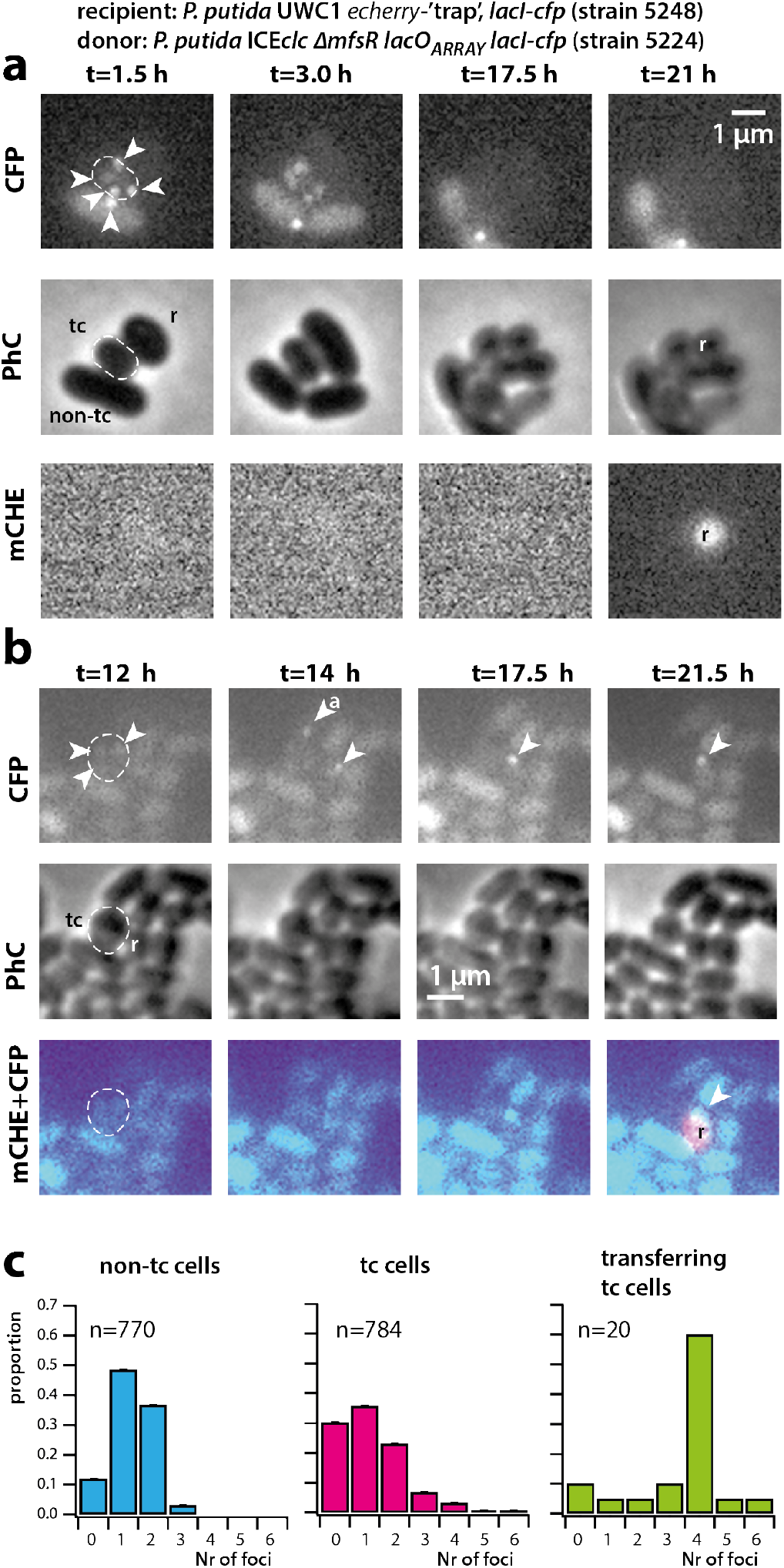
ICE transfer is favored from tc cells with higher copy number of excised ICE*clc*. **a, b**, Examples of ICE*clc* transfer from tc donor cells with excised and replicated ICE (note the 3–5 visible LacI-CFP foci in donor cells, stippled outlines) to neighbouring ICE-free recipient cells with the conditional trap (r) and appearance of mCherry fluorescence (mCHE) as a result of ICE-integration (t=21 h in **a** and t=21.5 h in **b**). Note how a LacI-CFP foci appears in each of two recipient cells in **b** (t=14 h), only one of which is integrated, whereas the other may be aborted (‘a’). **c**, observed foci distributions in tc donor cells with successful ICE*clc* transfer to recipient, compared to the foci distributions of all non-tc and tc cells of the same strain in absence of recipient.

## Discussion

It is increasingly recognized that selfish DNA elements mediating horizontal gene transfer have life-styles of their own, which are subject to adaptation and selection^3^. This is not only interesting as biological or molecular curiosity, but crucial to understand, given the role of such elements in promoting antibiotic resistances and xenometabolism in microbial communities^26^. Eventually, some elements may be more successful in distributing these genes than others, and some conditions may preferentially select for successful DNA-transferring elements. ICEs may be particularly relevant for this question as they have evolved exquisite regulatory systems to control their life-style, both in their vertical modes of co-replication with the bacterial chromosome, and the switch to horizontal transfer^1–3^. Although several different ICE models have contributed to the molecular understanding of ICE functioning, in particular studies on the ICE*clc* in *P. knackmussii* and *P. putida* have helped to elucidate the context of cellular differentiation and ecological fitness. As a result of spontaneous recent gene integration^27^, wild-type ICE*clc* is transferring at a sufficiently high rate (1 per 100) that single cell studies can be conducted, whereas most other wild-type ICE transfer at rates of 1 per 10^5^ or less (estimated in Ref^3^). Single cell studies revealed that ICE*clc* transfer is only accomplished from differentiated tc cells arising under no-growth (stationary phase) conditions^14,21^. The fitness strategy of the ICE is thus based on two pillars: vertical descent through co-replication with the chromosome and horizontal transfer via specialized tc cells. As cell division is perturbed in tc cells as a result of the ICE transfer process, too high transfer rates would compromise fitness of the host-ICE population of cells^14^. Too low transfer would not be in the fitness interest of the ICE, so that, depending on the host type and ICE model, some (dynamic) equilibrium arises that balances fitness loss in tc cells with ICE fitness gain from transfer and vertical descent^14^.

Interestingly, however, and this has been rarely recognized, the transient existence of tc cells necessitates ICE*clc* to transfer as optimally as possible in order to maximize horizontal fitness. The adaptations for this process can only be seen at individual cell level. Hence, we recognize for the first time that a transient phase of replication of excised ICE molecules in tc cells favors more effective transfer (Fig. 5C), which is a selectable feature increasing the ICE (horizontal) fitness. Mechanistically, this may occur through independent ICE transfer events at multiple positions around the cell, or through enhanced delivery rates of multiple single-stranded ICE-DNA at a single conjugative pore. Visual occurrence of quasi simultaneous multiple transfers from single donor to different neighbouring recipient cells (as in Fig. 5B) would favor the hypothesis of existence of multiple transfer pores, although these have so far not been seen.

We acknowledge that single molecule single cell studies carry many pitfalls, which we tried to control for as good as possible. The LacI-CFP system is not perfect and our data indicate that not all ICE copies might be detected by it. Therefore, it is difficult to obtain absolute discrimination between cells that lack the ICE or those with aberrant induction by arabinose, since both are characterized by the absence of LacI-CFP foci. Still, the high number of foci seen specifically in tc cells compared to non-tc is experimentally robust and statistically significant, and is consistent with the concept of replicating excised ICE*clc* in tc cells. The presence of two foci is explained by ongoing chromosome replication in dividing but not separated cells. Given the integration position of ICE*clc* in the *P. putida* chromosome close to the ter region (Fig. 3C), it is unlikely that renewed chromosome replication at the ori in dividing cells could produce 4 CFP foci. The observed three CFP foci in non-tc cells are therefore possibly the result of replication of integrated ICE*clc*, as was deduced and postulated for ICEBs1 in *Bacillus subtilis*^16,28,29^. Colabeling experiments showed that on average the TetR-YFP and LacI-CFP foci separate in tc cells, but not in non-tc cells. This indicates that the ICE molecule physically disengages from its nearby chromosomal location, and, consequently must be excised in tc cells. Most likely, therefore, even two CFP foci in small tc cells (< 1.8 µm) reflect excised ICE*clc*, and any number of foci equal to or larger than 3 in tc cells indicates replication of excised ICE*clc*. This conclusion is consistent with the absence of >3 foci in mutants without the *attL* recombination site, in which the ICE is chromosomally locked. Further consistent with population-based studies on other ICEs in *B. subtilis* and *Vibrio cholerae*^17,23^, the *traI* relaxase seems to be responsible for the replicative effect, since both *traI* and *ori1* deletions yielded tc cells with CFP foci numbers of less than 3. We thus feel relatively certain that foci numbers ≥ 3 in tc cells represent replicated excised ICE*clc* molecules.

Finally, although we cannot capture all possible ICE transfers, because many of them will go invisible, among those that we could capture (as in the experiments of Fig. 5), the majority of tc donor cells displayed multiple LacI-CFP foci, many more than expected by chance from the distribution of LacI-CFP foci seen among all tc cells (Fig. 5C). This does not exclude that cells with fewer LacI-CFP foci (and consequently, lower numbers of excised ICE-molecules) transfer at all, but indicates that tc cells with higher numbers of ICE-replicons have a higher chance of transferring the ICE. Transient replication of ICE upon excision is commonly interpreted as a selected feature to avoid ICE-loss in dividing cells, but we show that a higher number of ICE replicons directly translates into a higher rate or success of transfer, and therefore, gain of horizontal fitness.

## Methods

### Strains and culture conditions

*Escherichia coli* strains used for plasmid cloning were routinely cultured at 37°C in LB medium, while *P. putida* strains were grown at 30°C in LB or in 21C minimal medium (MM) (Gerhardt, 1981) supplemented with 5 mM 3-chlorobenzoate (3-CBA) or 10 mM Na-Succinate. Antibiotics were used at the following concentrations, if necessary: ampicillin (Amp) 100 µg ml^−1^, gentamicin (Gm) 20 µg ml^−1^, kanamycin (Km) 50-100 µg ml^−1^. Strains used in this study are listed in supplementary table S1.

### Strain constructions and DNA techniques

DNA manipulations and molecular techniques were performed according to standard procedures^30^ and recommendations by the reagent suppliers. Targeted chromosomal deletions and insertions in *P. putida* were created by recombination with non-replicating plasmid constructs and counter-selection techniques as previously described^20,31^. Recombination was facilitated by including regions up- and downstream to the targeted positions with sizes of approximately 0.7–kb.

To visualize and quantify ICE*clc*-containing DNA molecules in individual cells, we deployed the technique of *in-vivo* binding of fluorescently-labeled LacI or TetR to multiple tandem copies of their cognate binding sites^19^. A DNA fragment containing two times 120 *lacO* copies (*lacO_ARRAY_*), each interspaced by 10 random bp and with a Km-resistance gene in the middle^19^, was inserted within the *amnB* gene of ICE*clc*^8^ in *P. putida* (Fig. 1B). The *amnB* gene is part of a metabolic pathway involved in the degradation of 2-aminophenol and non-essential for the conjugative transfer of ICE*clc*^8^. The corresponding fragment was recovered from pLAU43^19^ using digestion with *Bam*HI and *Sal*I, and cloned into the *Pseudomonas* recombination vector pEMG^31^ (accession number JF965437), flanked by two 0.7-kb recombination fragments surrounding *amnB*. Similarly, a *tetO* array consisting of 2×120 *tetO* binding sites separated by a Gm-resistance gene was extracted from pLAU44^19^ using *Nhe*I and *Xba*I, cloned into pEMG and flanked with two fragments for recombination next to the gene Pp_1867, which is located 12 kb upstream of the insertion site of ICE*clc* in the genome of *P. putida* (accession number NC_002947.4). Proper recombination and marker insertion was verified by PCR amplification and sequencing.

The *araC, lacI-cfp* fragment used for arabinose-inducible expression of LacI-CFP under control of AraC was amplified from pLAU53^19^, verified by sequencing and cloned in a mini-Tn*7* delivery plasmid^32^ (accession number AY599231) using *Spe*I and *Hind*III. The *araC, lacI-cfp, tetR-yfp* fragment used for ectopic expression of both LacI-CFP and TetR-YFP was retrieved from pLAU53 using digestion with *Sgr*AI and *Hind*III, and inserted into the mini-T*n*7 delivery plasmid at *Xma*I and *Hind*III positions. The resulting plasmids were co-transformed with the Tn7-expressing helper plasmid pUX-BF13^33^ into the different *P. putida* strains (Supplementary Table S1). After selection of transformants for the respective antibiotic resistance markers expressed by the mini-Tn*7* cassette, its proper site-specific insertion at the *glmS* site was verified by PCR amplification.

To differentiate non-tc and tc cells we used the ICE tc-cell specific P_inR_-promoter^13,20^, which was fused to a promoterless *echerry* gene in a transcriptionally shielded mini-Tn*5* transposon, and integrated in single copy on the *P. putida* chromosome using a mini-Tn*5* delivery vector, as previously described^14^. Three independent mini-Tn*5* insertions were kept for each derivative strain.

### Epifluorescence microscopy

Fluorescent protein expression in individual cells was examined by epifluorescence microscopy on a Nikon Inverted Microscope Eclipse Ti-E, equipped with a Perfect Focus System (PFS), pE-100 CoolLED and a Plan Apo l 100 ×1.45 Oil objective (Nikon), installed in a controlled temperature room (22 °C).

Cell growth and tc cell development were followed in 50-h long time-lapse experiments, with cells seeded on round (ø 1 cm diameter, 1 mm thick) 1% agarose disks placed inside closed sterilized metal microscope chambers^14,31^. Surfaces were inoculated with late stationary phase (96 h) precultures, to ensure the presence of tc cells at the beginning of the experiment. Precultures were prepared by transferring 100 µl of an over-night grown culture on LB with antibiotics to maintain selection of the chromosomal markers into an Erlenmeyer flask containing 20 ml of MM with 5 mM CBA (without antibiotics). This culture was incubated for 96 h at 30°C with 200 rpm rotary shaking (cells reach stationary phase after 24 h). If relevant, at this point L-arabinose was added to the culture at a final concentration of 100 mg l^−1^ to induce expression of *lacI-cfp* from P_BAD_. After 90 min incubation, 1 ml of the culture was centrifuged for 2 min at 18,000 × *g* to collect the cells, which were re-suspended in 10 ml MM without added carbon substrate. 6 µl of this washed preculture was then spreaded per agarose disk, which further containing 0.1 mM 3CBA in MM and 10 mg l^−1^ L-arabinose to maintain induction from P_BAD_.

For observation of chromosome replication in dividing *P. putida* cells with both LacI-CFP and TetR-YFP labeling (strain 5601), we imaged cells directly (i.e., without time-lapse) from a liquid culture in MM with 5 mM 3CBA and 10 mg l^−1^ L-arabinose incubated for 4 h at 30°C. For imaging, cells were concentrated and resuspended as described above, and spread on 1% agarose surface on microscope slides. This culture was prepared by tenfold dilution from a preculture in MM with 5 mM 3CBA that had been grown to stationary phase for 48 h (to ensure tc cell arisal), after which 100 mg l^−1^ L-arabinose had been added for 90 min to express LacI-CFP and TetR-YFP. An incubation of 4 h is sufficient to revive the cells from stationary phase and resume cell division in tc and non-tc cells (note that any tc cells lysing within this period will be lost from the analysis).

In case of ICE*clc* time-lapse transfer experiments, donor cells were prepared as described above. The *P. putida* recipient strain with the conditional eCherry-fluorescent ICE-integration trap and ectopically expressing LacI-CFP (strain 5248) was grown with 10 mM succinate for 24 h, and incubated with 100 mg l^−1^ L-arabinose for 90 min. Donor and recipient cells were washed as described above, resuspended in MM without carbon source, mixed in a 1:2 (*v*/*v*) ratio, respectively, and seeded on 1% agarose disk surface with 0.1 mM 3CBA and 10 mg L^−1^ L-arabinose as previously for donor cells alone.

Seeded agarose disks were turned upside down, cells facing the lower coverslip, and enclosed in an autoclaved microscopy chamber (Perfusion Chamber, H. Saur Laborbedarf, Germany). Assembled chambers contained four simultaneous patches, one of which remained non-inoculated and served to pause the microscope objective in between imaging and avoid light–induced stress on the cells. Chambers were adapted for 1 h to temperature (22 °C) and humidity of the microscope room, before starting the time-lapse experiment. Images were recorded at a light intensity of 10% (Solar light engine, LED power 4%) and an ORCA-flash 4.0 Camera (Hamamatsu). Exposure times for phase-contrast images were 50 or 100 ms, for mCherry fluorescence 20 or 50 ms; for CFP fluorescence 200 or 250 ms, and for YFP fluorescence 250 ms. Four positions on each disk were programmed in MicroManager (version 1.4.22), and were imaged every 30 min during 50 h.

### Image analysis

Fluorescence values of single cells obtained from snapshot microscopy experiments were extracted using an in-house Matlab script as described previously^14^ and subpopulations were quantified from quantile-quantile plotting^21^. tc cells were categorized on the basis of eCherry-fluorescence expressed from the single copy P_inR_-promoter in quantile-quantile analysis, which scores the deviation of the observed distribution of eCherry fluorescence among individual cells to the expected normal distribution assuming a single population^12,21^.

Individual cells in time-lapse image series (up to 100 frames) were segmented using SuperSegger^34^, and both cellular fluorescence as well as the fluorescence intensities, scores and positions of up to 9 foci in individual cells were extracted. Optimized segmentation constants for *P. putida* were derived from the *training your own constants* subprocedure in SuperSegger. Extracted data were then analyzed with a homemade Matlab script (Supplementary file S1), to (i) identify tc and non-tc cells, (ii) to derive the genealogy of all cells and link them within growing microcolonies (cell ID, frame of birth, frame of death, mother and daugther IDs), and (iii) to count the position, number, normalized fluorescence intensity and scores of individual cell foci over time. tc cells were identified in the first image frame on the basis of quantile-quantile plotting of eCherry fluorescence values as the subpopulation with the highest eCherry fluorescence, whereas the largest subpopulation with the lower average eCherry fluorescence was considered to contain non-tc cells. Mother cells forming microcolonies of less than 8 cells were further excluded of the group of non-tc cells, since they may consist of tc cells with low eCherry starting values. This procedure was justified based on previous observations of poor re-growth of tc cells^12^. Individual cells with more than 4 foci were examined manually using the superSeggerViewerGUI mode of SuperSegger, and incorrectly segmented cells were removed from the final analysis. Foci distributions among the groups of tc and non-tc cells were normalized to within-group percentages and compared using the Fisher’s exact test, given the absence of an *a priori* distribution function.

## Acknowledgements

The authors thank Stephan Gruber and Nicolas Carraro for critical reading of the manuscript. We thank Stella Stylianidou for her help and advice with SuperSegger. This work was supported by the Swiss National Science Foundation grants 310030B_156926 and 31003A_175638.

## Author Contributions

F.D., R.M. and J.v.d.M. designed experiments. F.D. and R. M. performed experiments. F.D., R.M. and J.v.d.M. analyzed data. F.D., R.M. and J.v.d.M. wrote the manuscript.

## Additional Information

Supplementary information is available for this paper at xxx

Correspondence and requests for materials should be addressed to J.v.d.M.

